# Glioblastoma extracellular vesicles influence glial cell hyaluronic acid deposition to promote invasiveness

**DOI:** 10.1101/2022.02.11.480036

**Authors:** Dominik Koessinger, David Novo, Anna Koessinger, America Campos, Jasmine Peters, Louise Dutton, Peggy Paschke, Désirée Zerbst, Madeleine Moore, Louise Mitchell, Matthew Neilson, Katrina Stevenson, Anthony Chalmers, Stephen Tait, Joanna Birch, Jim Norman

**Affiliations:** Cancer Research UK Beatson Institute, Glasgow, G61 1BD, UK; School of Cancer Sciences, University of Glasgow, Glasgow G61 1QH, U.K; Department of Neurosurgery, Freiburg University Hospital, Breisacher Str. 64, 79106 Freiburg, Germany; Current address: Francis Crick Institute, 1 Midland Road, London NW1 1ST, UK; Lead contact authors; Shared first authorship

**Keywords:** extracellular vesicles, GBM, invasion, astrocytes, extracellular matrix.

## Abstract

**Background:** Infiltration of glioblastoma (GBM) throughout the brain leads to its inevitable recurrence following standard-of-care treatments, such as surgical resection, chemo- and radio-therapy. A deeper understanding of the mechanisms invoked by GMB to infiltrate the brain is needed to develop approaches to contain the disease and reduce recurrence. The aim of this study was to discover mechanisms through which extracellular vesicles (EVs) released by GBM influence the brain microenvironment to facilitate infiltration, and to determine how altered extracellular matrix (ECM) deposition by glial cells might contribute to this.

**Methods:** CRISPR was used to delete genes, previously established to drive carcinoma invasiveness and EV production, from patient-derived primary and GBM cell lines. We purified and characterised EVs released by these cells, assessed their capacity to foster pro-migratory microenvironments in mouse brain slices, and evaluated the contribution made by astrocyte-derived extracellular matrix (ECM) to this. Finally, we determined how CRISPR-mediated deletion of genes, which we had found to control EV-mediated communication between GBM cells and astrocytes, influenced GBM infiltration when orthotopically injected into CD1-nude mice.

**Results:** GBM cells expressing a p53 mutant (p53^273H^) with established pro-invasive gain-of-function release EVs containing a sialomucin, podocalyxin (PODXL), which encourages astrocytes to deposit ECM with increased levels of hyaluronic acid (HA). This HA-rich ECM, in turn, promotes migration of GBM cells. Consistently, CRISPR-mediated deletion of *PODXL* opposes infiltration of GBM *in vivo*.

**Conclusions:** This work describes several key components of an EV-mediated mechanism though which GBM cells educate astrocytes to support infiltration of the surrounding healthy brain tissue.

**KEY POINTS:** The p53^R273H^ oncogene encourages GBM cells to release EVs containing podocalyxin. Podocalyxin-containing EVs from GBM increase hyaluronic acid production by astrocytes. Hyaluronic acid production by astrocytes drives GBM migration.

**IMPORTANCE OF THE STUDY:** The infiltrative behaviour of glioblastoma (GBM) leads to widespread dissemination of cancer cells throughout the brain. Thus, even following successful resection of the primary tumour these disseminated cells inevitably contribute to post-surgical relapse. In this study, we have discovered a new mechanism through which GBM can release small extracellular vesicles (EVs) to reprogramme extracellular matrix (ECM) production by astrocytes in a way that supports increased invasive behaviour of the GBM cells. Moreover, we have discovered several key components of the pathway which contribute to this EV-mediated GBM-glial cell communication. Principal amongst these, we show that a particular mutant of the p53 tumour suppressor, p53^273H^ drives the release of EVs which foster the deposition of pro-invasive ECM by astrocytes. This study provides mechanistic insight into why brain tumours expressing p53^273H^ are associated with particularly poor patient survival and highlights the possibility of deploying agents which target astrocyte ECM deposition to reduce the morbidity of p53^273H^- expressing GBM.

## INTRODUCTION

Glioblastoma (GBM) exhibit invasive/infiltrative behaviour which is largely responsible for the intractable nature of the disease ^1^. At diagnosis GBM usually display widespread infiltration of the surrounding brain tissue which precludes *in toto* resection and focussed radiotherapy and, therefore, leads to relapse. Furthermore, widespread insinuation of invasive cells into brain tissue is responsible for many of the neurological and cognitive symptoms which contribute to GBM’s morbidity ^2^. Thus, an understanding of the mechanisms though which GBM acquire infiltrative/invasive characteristics, and the cellular mechanisms involved in this, is necessary to assist development of pharmacological strategies to control the disease following resection both before and following chemo and radiotherapy – thus improving accessibility for treatment in the case of relapse and ameliorating its associated neurological and cognitive dysfunction.

The ways in which cancers influence the microenvironment in distant organs to prime these for metastatic colonisation are now becoming clear. Soluble components of the cancer secretome can mobilise myeloid cell populations to metastatic target organs to generate immunosuppressive microenvironments ^3^. Additionally, tumour-derived extracellular vesicles (EVs) can prime metastatic niches via alterations to the vasculature and recruitment of myeloid cells with immunosuppressive phenotypes. Furthermore, tumour EVs can alter the extracellular matrix (ECM) within organs such as the liver and lung to generate pro-invasive microenvironments which favour metastatic seeding. ^4,5^

Infiltrative behaviour in GBM can be mediated via both cell autonomous and complex intercellular communication processes involving alterations in the nearby and distant brain microenvironment. Invading glioma cells produce a range of protrusion-types – including tumour microtubes and tunnelling nanotubes – to form functional intercellular communication networks which alter the local and wider brain microenvironment to favour infiltration and therapy resistance ^6,7^. EVs also mediate communication between GBM cells and the brain microenvironment in a way that would be expected to favour tumour cell proliferation, angiogenesis and immune infiltration ^8,9^. As with metastatic priming, a number of studies now indicate that GBM EVs increase recruitment of myeloid cell populations with tumour promoting and/or immunosuppressive properties ^10-13^. However, despite the importance of the ECM in GBM infiltration, it is currently unclear how GBM EVs might alter the brain ECM, and what the role of the brain’s main ECM depositing cell-type, the astrocyte, might be in this regard. Therefore, we have studied how an oncogene – the p53^273H^ mutant – which is closely associated with poor clinical outcomes - alters the constitution of EVs released by patient-derived glioma cells. We report that EVs released by p53^273H^-expressing GBM cells are responsible for enabling their invasive behaviour *in vivo*, and this is due to the ability of these EVs to influence the hyaluronic acid (HA) content of ECM deposited by astrocytes.

## MATERIALS AND METHODS

### Patient-derived GBM cells

The E2 and G7 GBM cell lines were obtained from Colin Watts (Cambridge, now Birmingham) and chosen for their characteristic growth pattern in CD1 nude mice xenografts ^14^. Cells were originally isolated from fresh tumour tissue from anonymised patients diagnosed with glioblastoma and continued expression of glioblastoma-characteristic cellular markers was compared to previous publications using these cell lines ^15-18^. GBM cells were cultured on Matrigel (diluted 1:40) under serum-free conditions in adDMEM/F12, 1% B27, 0.5% N2, 4μg/ml heparin, 10ng/ml fibroblast growth factor 2 (bFGF), 20 ng/ml epidermal growth factor (EGF) and 1% L-glutamine. The U373 GBM cell line was cultured in DMEM containing 10% foetal calf serum on uncoated plastic surfaces.

### CRISPR and transfections

For CRISPR/cas9-mediated gene knockouts, the following guide sequences were cloned into the lentiCRISPR vector (Addgene plasmid #52961 – deposited by the Zhang lab ^19^):

TP53 #1 forward CACCGCGCTATCTGAGCAGCGCTCA

TP53 #1 reverse AAACTGAGCGCTGCTCAGATAGCGC

TP53 #2 forward CACCGCCCCGGACGATATTGAACAA

TP53 #2 reverse AAACTTGTTCAATATCGTCCGGGGC

PODXL #1 forward CACCGCAGCTCGTCCTGAACCTCAC

PODXL #1 reverse AAACGTGAGGTTCAGGACGAGCTGC

PODXL #2 forward CACCGGGTGTTCTCAATGCCGTTGC

PODXL #2 reverse AAACGCAACGGCATTGAGAACACCC

Rab35 #1 forward CACCGCTTGAAATCCACTCCGATCG

Rab35 #1 reverse AAACCGATCGGAGTGGATTTCAAGC

Rab35 #2 forward CACCGGAAGATGCCTACAAATTCGC

Rab35 #2 reverse AAACGCGAATTTGTAGGCATCTTCC

Active lentiviruses were produced using HEK293FT cells as the packing line. E2 GBM cells were plated onto Matrigel, transduced with lentiviruses and selected using puromyucin (1 μg/ml) or blasticidin (5 μg/ml).

For overexpression of PODXL, the sequence for human PODXL (hPODXL) was cloned into the pQCXlZ-eGFP-C1 retroviral vector (a gift from David Bryant) and Phoenix-Ampho cells were used as the host packaging line. E2 GBM cells were plated onto Matrigel, transduced with retroviruses and selected using Zeocin (1 mg/ml).

### EV purification and nanoparticle tracking analysis

EVs were collected via differential centrifugation of cell-conditioned medium as described previously ^5,20^. Nanoparticle tracking analysis was carried out using the NanoSight LM10 instrument (Malvern Panalytical) according to the manufacturer’s instructions. To measure uptake by astrocytes, purified EVs were labelled by incubation with PHK67 (2μM) for 5 min. Excess dye was removed by ultracentrifugation (100,000 g for 70 min) and labelled EVs were added to primary cultured astrocytes for 24hr. Recipient astrocytes were then analysed using flow cytometry.

### ECM generation

Primary rat astrocytes were grown to 80% confluence in 15cm dishes before being incubated in the presence or absence of EVs at a concentration of 1 × 10^9^ particles/ml for 72 hr. EV-treated astrocytes were re-plated at 1 × 10^6^ cells/well into 6-well dishes pre-coated with 0.2% gelatin, which was subsequently crosslinked with 1% glutaraldehyde for 30 min, quenched in 1M glycine for 20min. Astrocytes were allowed to deposit ECM for 6 days in medium supplemented with 50 μg/mL ascorbic acid in the presence or absence of the DGK-inhibitor R59022 (10μM) (Sigma) or hyaluronidase (Hase) (50 µg/mL; Type I-S, Sigma H3506) where indicated. ECM was then de-cellularised by incubation with PBS containing 20mM NH_4_OH and 0.5% Triton X-100.

### Immunofluorescence detection of HA and CSPG in astrocyte-deposited ECM

To visualise HA, samples were incubated with biotin-conjugated hyaluronic acid binding protein (HABP) (Sigma 385911), followed by streptavidin conjugated to AlexaFluor-488 (ThermoFisher S11223), and for chondroitin sulphate proteoglycan (CSPG), a mouse monoclonal antibody (Sigma C8035) was used followed by an anti-mouse secondary antibody conjugated to AlexaFluor-488. Visualised was by confocal microscopy using an Olympus Fluoview FV1000 microscope. Z-stacks were acquired from the substrate to the upper surface of the cultures at intervals of 0.5 μm.

### GBM cell migration

GBM cells were seeded onto de-cellularised ECMs or coronal slices from DGKα^+/+^ or DGKα^-/-^ 5-8 week female C57Bl/6 mice. GFP-expressing cells were visualised using a Nikon A1R microscope with frames being captured every 10 min for 16 hr.

### Orthotopic xenografts

Animal experiments were performed under the relevant home office licence (project licence PPL P4A277133) and in accordance with ARRIVE guidelines. All experiments had ethical approval from the University of Glasgow under the Animal (Scientific Procedures) Act 1986 and the EU directive 2010. 7 week old, female CD1-nude mice (Charles River) were anesthetised using isoflurane and placed in a stereotactic frame. To prevent eye desiccation, Lacrilube eye cream was applied. For analgesia, diluted Veterigesic was injected subcutaneously at a final dose of 20 μg/kg. The skin of the surgical area was disinfected using Hydrex skin disinfectant. The mouse was then covered using sterile drapes. Prior to incision, anaesthesia depth was assessed via pedal response. Subsequently the skin was incised along the sagittal suture and periosteum was removed using sterile cotton swabs. A burr hole craniotomy was performed using an electric hand drill 3 mm rostral and 2 mm lateral of the bregma over the right hemisphere. Subsequently, 1 × 10^5^ GBM cells were injected (0.2 × 10^5^ cells/μl in PBS at a rate of 2 μl/min for 2.5 min) using a Hamilton syringe inserted 3mm into the brain. Skin was adapted and Vetbond tissue glue applied. Mice were put in pre-warmed recovery cages and continuously monitored until mobile. For postoperative analgesia, mice were provided with Rimadyl-containing drinking water for 48 hr postoperatively. Mice were continuously monitored for throughout the course of the experiment, and humanely sacrificed either upon display of neurological (such as hemiparesis or paraplegia) or general (hunched posture, reduced mobility, and/or weight loss >20%) symptoms, or timed end-point of 9 weeks following engraftment of GBM cells. Formalin fixed, 4μm thick coronal mouse brain sections were stained for Ki67 as previously described ^14^.

## RESULTS

### EVs from infiltrative GBM cells foster a pro-migratory microenvironment in brain slices

Two primary patient-derived glioma stem-like cell (GSC) lines, G7 and E2, derived from resected patient tissue^15,18^ were chosen in view of the markedly different invasive/infiltrative characteristics that they display *in vivo*. Indeed, G7 GSC grow as a solid tumour with moderately invasive margins, whereas E2 disseminate throughout the brain as widely scattered tumour cells ^14^. To determine whether this was owing to their intrinsically different migratory behaviour, we plated GFP-expressing G7 and E2 cells onto brain slices from 5–8-week Cl57/Bl6 mice and recorded their movement using fluorescence time-lapse microscopy. Surprisingly, both the G7 and E2 cell lines migrated similarly (and poorly) on brain slices (Figure S1A) despite their markedly different invasive behaviour *in vivo* ^14^. We hypothesised that the secretome of GBM cells may induce changes in the brain microenvironment which promote invasiveness, and that this may take some time to establish. As EVs are a component of the tumour secretome with key roles in influencing microenvironments both locally and at distance from the primary tumour, we purified EVs from GSC-exposed medium and analysed these using nanoparticle tracking. G7 and E2 cells released EVs in similar quantities and with indistinguishable size distributions (Figure S1B). We then incubated brain slices with EVs from G7 or E2 cells for 72 hr, subsequently plated GFP-expressing E2 or G7 cells onto these and measured their migration. This indicated that pre-treatment of brain slices with EVs from E2 (but not G7) cells increased migration of subsequently-plated GBM cells (Figure 1A). Consistently, GBM cells plated onto brain slices pre-treated with EVs from E2 cells displayed a more invasive phenotype characterised by extension of invasive protrusions which altered the cell shape (Figure 1B).

**Figure 1:**
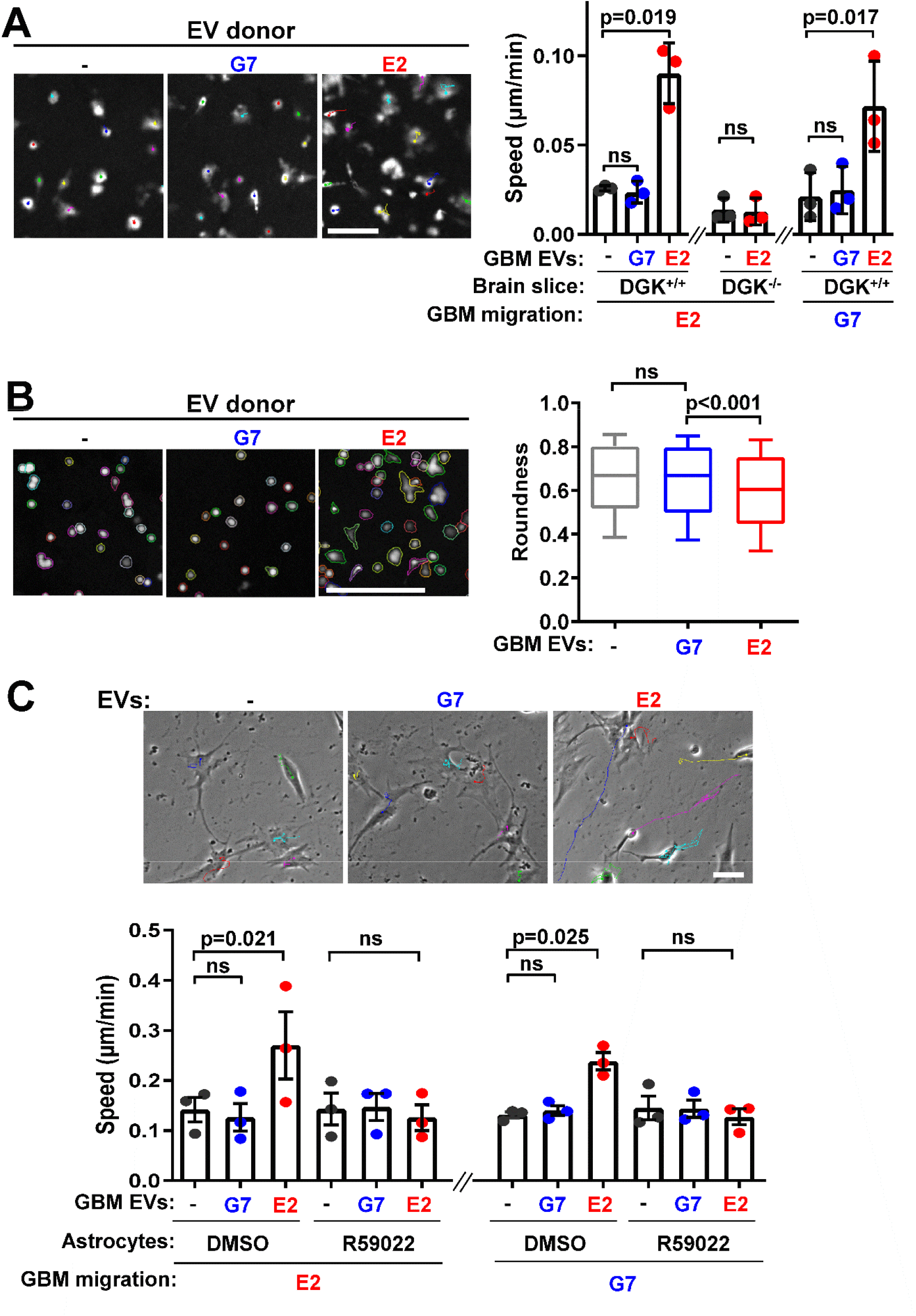
EVs from infiltrative GMB cells foster pro-migratory microenvironments. **A, B)** Brain slices from DGK^+/+^ or DGK^-/-^ mice were treated for 72 hr with equal quantities of EVs from E2 or G7 patient-derived glioma stem-like (GBM) cells or left untreated (-). GFP-expressing E2 or G7 cells were then plated onto slices and migration speed (A) and shape (B) determined. In (A) values are mean ± sem (n=3 independent experiments, paired t-test, 60 cell-tracks/condition/experiment). In (B) bars are median, boxes denote interquartile range and whiskers the 10 – 90 % range. (n=3, unpaired t-test). Bars are 50 μm. **C)** Astrocytes were treated for 72 hr with EVs from E2 or G7 cells or were left untreated (-). Astrocytes were then allowed to deposit ECM for 6 days in the presence or absence of 10 μM R59022 or vehicle control (DMSO). ECM was decellularized and E2 (left graph and micrographs) or G7 (right graph) cells plated onto these. GBM cell migration was then determined using time-lapse microscopy. Values are mean ± sem (n=3 independent experiments, paired t-test, 60 cell-tracks/condition/experiment).

The lipid kinase, diacylglycerol kinase-α (DGKα) must be expressed in fibroblasts for them to generate pro-invasive microenvironments in response to EVs from primary tumours^5^. We deployed DGKα knockout (DGKα^-/-^) mice to determine whether this kinase is required for a ‘recipient’ brain cell population to generate a pro-migratory microenvironment in response to GBM EVs. Brain slices from 5-8 week DGKα^-/-^ Cl57/Bl6 mice were treated with EVs from E2 cells for 72 hr, and GFP-expressing E2 cells subsequently plated onto these. GBM cells migrated poorly on brain slices from DGKα^-/-^ mice (as they did on slices from DGK^+/+^ mice) and this was unaffected by pre-treatment with EVs from E2 cells, indicating that expression of DGKα is required for a brain cell population to generate a pro-migratory microenvironment in response to GBM EVs (Figure 1A).

### EVs from infiltrative GBM cells encourage astrocytes to deposit pro-migratory ECM

A principal task of astrocytes is to deposit ECM in the brain and thus maintain its appropriate microenvironment. Experiments in which we labelled EVs with fluorescent dyes and incubated them with primary cultured astrocytes indicated that these glial cells are capable of assimilating and, therefore, potentially responding to, EVs from both G7 and E2 GBM cells (Fig. S2). We, therefore, incubated primary -cultured astrocytes with EVs from E2 or G7 cells for 72 hours, re-plated the astrocytes and allowed them to deposit ECM for 6 days. Astrocyte cultures were then de-cellularised and migration of GBM cells on these ECMs was determined using time-lapse microscopy. Pre-treatment of astrocytes with EVs from E2 (but not G7) cells prior to ECM generation significantly increased the speed of both E2 and G7 GBM cells plated on these matrices (Figure 1C). Moreover, addition of a DGKα inhibitor (R59022) ^21,22^ to the astrocyte cultures opposed the ability of EVs from E2 cells to encourage astrocytes to deposit ECM with increased capacity to support GBM cell migration.

### Deletion of p53^273H^ reduces the ability of EVs from GBM cells to foster a pro-migratory microenvironment

An analysis of deep-sequencing data revealed that the highly infiltrative E2 cells express the p53^R273H^ mutant (83% of reads), which has well-characterised pro-invasive/metastatic gain-of-function properties ^23-26^ (Supplementary table S1A). Contrastingly, expression of mutated p53s in the less invasive G7 cells is less consistent, with reads being detected for various mutations (p.R248Q p.R282W p72r, each approx. 50% of reads) (Supplementary table S1B) for which a connection to invasion and metastasis is not established ^27,28^. To determine whether expression of p53^273H^ contributes to the generation of pro-invasive/migratory microenvironments, we generated clones of E2 cells in which the gene for p53 was targeted using two independent CRISPR guide sequences. Western blotting confirmed deletion of p53^273H^ and that expression of neuronal stem cell markers, Nestin and SOX2, had not been compromised during this procedure (Figure 2A). Moreover, deletion of mutant p53 did not alter proliferation (Figure 2B), nor the quantity or size-distribution of EVs released from E2 cells (Figure 2C). As previous work indicates that mutant p53 generates pro-invasive/migratory niches by modulating the quantity of the sialomucin, podocalyxin (PODXL) sorted into EVs ^5^, we determined the influence of p53^273H^ on PODXL levels in EVs from E2 cells. PODXL levels in EVs were increased by deletion of mutant p53^273H^ (Figure 2D); an observation consistent with previous findings in carcinoma cells, and which moots the probability that EVs from p53-KO E2 cells might have altered ability to foster pro-invasive microenvironments ^5^. Indeed, EVs from p53-KO E2 cells displayed reduced capacity to influence astrocyte ECM deposition in a way that supported migration of GBM cells (Figure 2E). To confirm p53^R273H^’s role in influencing the ability of EVs to foster deposition of pro-migratory ECM, we deployed another GBM cell line, U373 - which bears this mutation ^29^ - and generated clones in which the mutant p53 was targeted using two independent CRISPR guides (Fig. 2A). Deletion of p53^R273H^ did not alter U373 cell proliferation, nor the quantity of EVs that they release (Fig. 2B, C). Nevertheless, in a similar manner to that observed for E2 cells, deletion of p53^R273H^ increased the PODXL content of EVs released by U373 cells (Fig. 2D) and, consistently, reduced their capacity to persuade astrocytes to deposit pro-migratory ECM (Fig. 2E). Furthermore, addition of a DGKα inhibitor (R59022) to the astrocyte cultures opposed the ability of EVs from both control U373 and E2 cells to evoke deposition of pro-migratory ECM by astrocytes. Finally, treatment of astrocytes with R59022 did not further reduce the (already compromised) migration-supporting properties of ECM deposited by astrocytes treated with EVs from p52^R273H^ CRISPR E2 or U373 cells (Fig. 2E).

**Figure 2:**
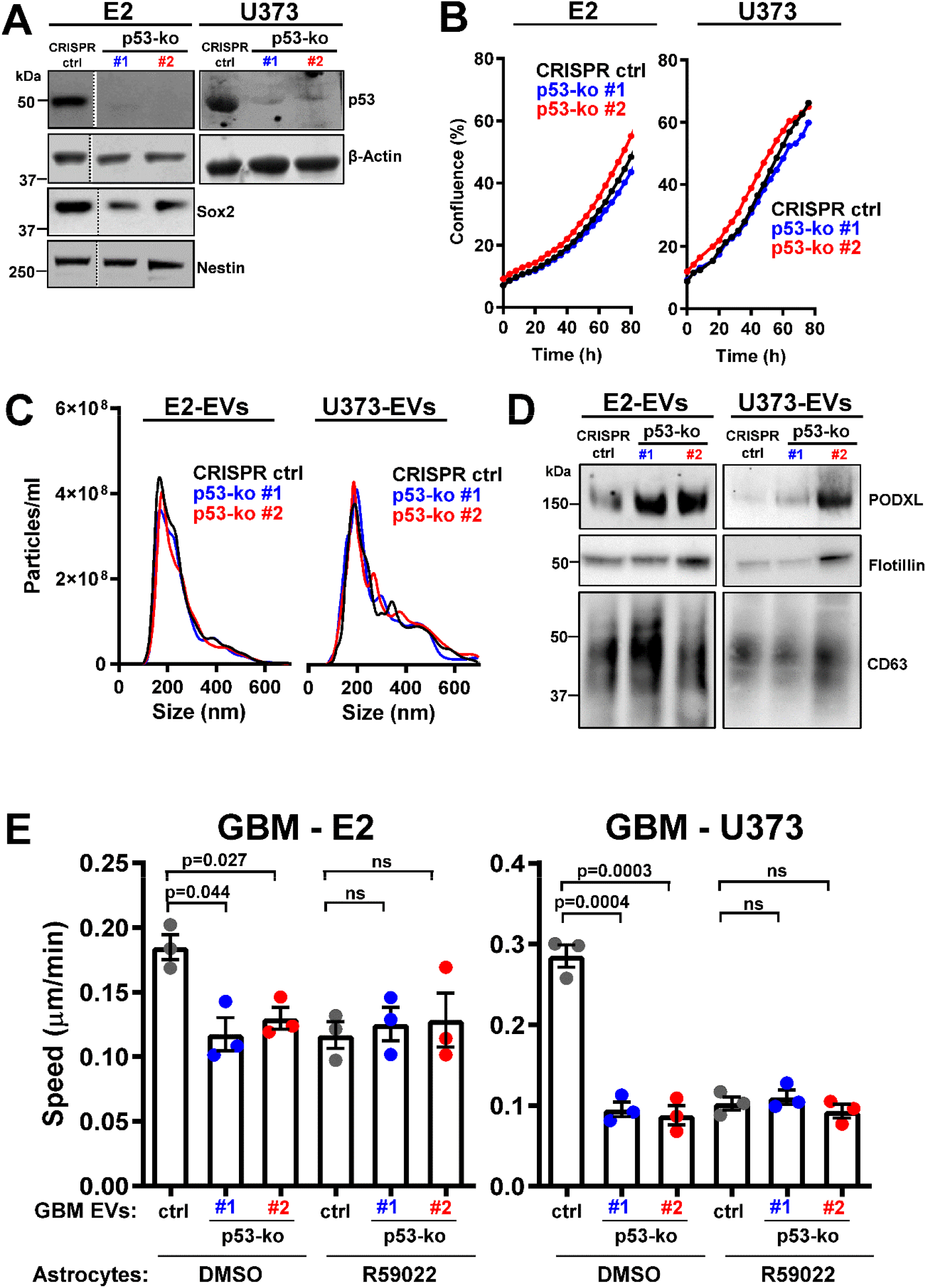
Deletion of p53^273H^ abrogates the ability of GBM cells to produce EVs which encourage astrocytes to deposit pro-migratory ECM. **(A-B)** Characterisation of control (CRISPR-ctrl) and p53 knockout (p53-ko #1 and #2) E2 and U373 cells: **(A)** confirms deletion of p53^273H^ in E2 and U373 cells, and expression of stem cell markers, nestin and SOX2 in E2 cells; **(B)** indicates that deletion of mutant p53 does not influence cell growth in E2 or U373 cells. **(C-D)** Characterisation of EV release by control and p53 knockout E2 and U373 cells: The number and size-distribution of EVs from CRISPR-ctrl and p53-ko cells was analysed using nanoparticle tracking **(C)**, and their PODXL, flotillin and CD63 content was determined by Western blot **(D)**. **(E)** Astrocytes were treated for 72 hr with EVs from control and p53 knockout E2 or U373 cells. Astrocytes were then allowed to deposit ECM for 6 days in the presence or absence of R59022, and migration of E2 cells on these ECMs determined as for Fig. 1C. Values are mean ± sem (n=3 independent experiments, paired t-test, 60 cell-tracks/condition/experiment).

### PODXL content of GBM EVs is critical to influencing astrocyte ECM deposition

The PODXL content of carcinoma cell EVs must be within a certain range for them to encourage fibroblasts to deposit pro-migratory ECM, and expression of mutant p53 functions to move EV PODXL levels into this range ^5^. To determine whether PODXL levels in EVs from GBM is causally linked to their ability to alter ECM deposition by astrocytes, we generated E2 cells in which PODXL levels were either increased (by over-expression) or reduced (using CRISPR). Furthermore, because Rab35 controls PODXL trafficking to EVs ^5^, we also generated Rab35 knockout GBM cells. Immunoblotting confirmed that EV PODXL levels were respectively increased or decreased by PODXL over-expression or knockout (Figure 3A, B) and deletion of Rab35 reduced EV PODXL levels (Figure 3C). These manipulations of PODXL and/or Rab35 levels did not alter GBM cell proliferation nor expression of neuronal stem cell markers (Figure 3A-C). We then pre-treated astrocytes with EVs from PODXL over-expressing or -deficient, and Rab35 knockout E2 cells for 72h, and allowed them to deposit ECM for a further 5 days. Astrocyte-derived ECM was de-cellularised, and migration of GBM cells on these ECMs determined using time-lapse microscopy. EVs from PODXL overexpressing (Figure 4A) and from PODXL (Figure 4B) or Rab35 (Figure 4C) knockout E2 cells were unable to influence astrocyte ECM deposition in a way that supported increased migration of GBM cells. Finally, we pre-treated mouse brain slices with EVs from control and PODXL or Rab35 knockout E2 cells and subsequently plated GBM cells onto these. This indicated that knockout of PODXL or Rab35 opposed the ability of EVs from GBM cells to foster pro-migratory microenvironments in the brain (Figure 4D). These data indicate that expression of p53^273H^ in GBM cells fosters a pro-invasive brain microenvironment, and that this is via regulation of EV PODXL content and the influence of this on the ECM depositing capacity of astrocytes.

**Figure 3:**
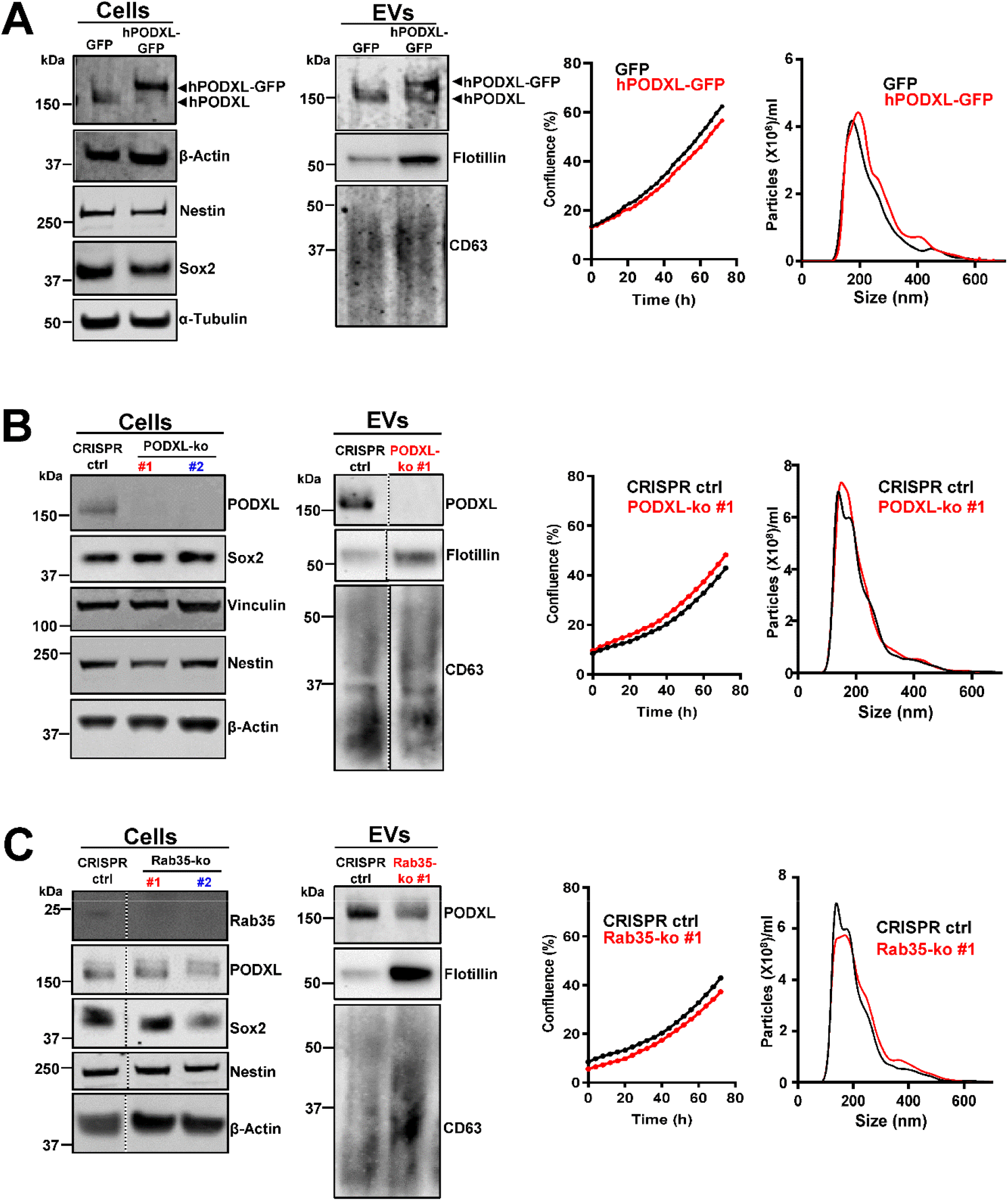
Manipulation of PODXL content in GBM EVs. E2 GBM cells were stably transfected with lentiviral vectors encoding GFP-tagged PODXL (hPODXL-GFP) or GFP **(A)**, or with CRISPR/Cas9 vectors containing 2 independent guide sequences targeting PODXL (PODXL-ko #1 and #2) **(B)**, Rab35 (Rab35-ko#1 and #2) **(C)** or empty vector control (CRISPR-ctrl). PODXL overexpression, deletion of PODXL and Rab35, expression of stem cell markers, nestin and SOX2 were confirmed by Western blotting. PODXL, flotillin and CD63 content of EVs was also determined by Western blotting. Cell proliferation and the number and size-distribution of EV were determined as for Fig. 2B & C.

**Figure 4:**
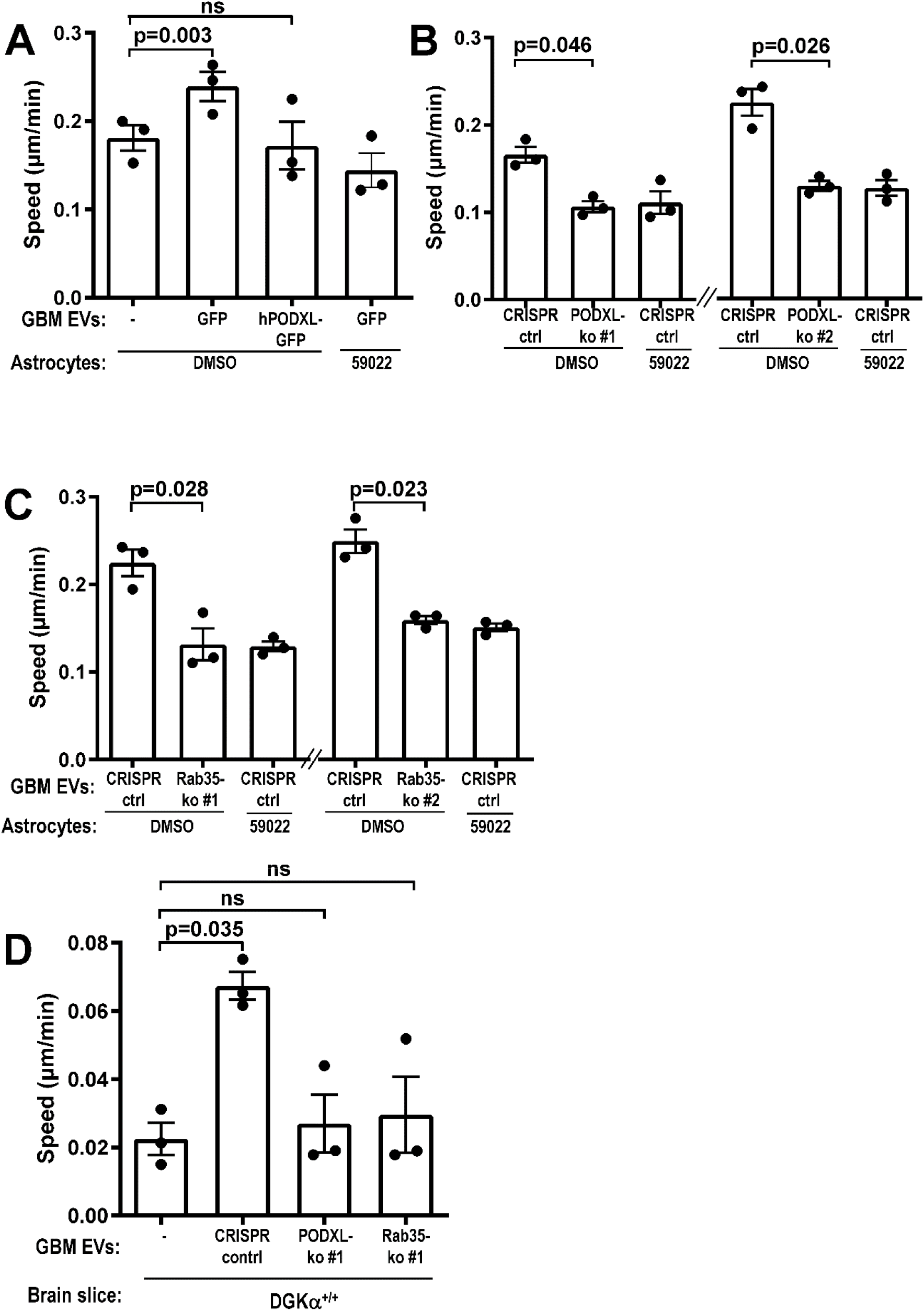
The level of PODXL in GBM Evs is critical for production of pro-migratory ECM by astrocytes. Astrocytes **(A-C)** or brain slices **(D)** were treated with EVs from GBM cells in which PODXL or Rab35 levels were manipulated (see Fig. 3). Astrocytes were then allowed to deposit ECM for 6 days in the presence or absence of R59022 or vehicle control (DMSO), and migration of E2 cells on these ECMs determined as for Fig. 1C. Values are mean ± sem (n=3 independent experiments, paired t-test, 60 cell-tracks/condition/experiment).

### EVs promote GBM cell migration by influencing the hyaluronic acid content of astrocyte ECM

EVs from mutant p53-expressing carcinoma cells generate pro-invasive/migratory niches in metastatic target organs (such as the lung), by influencing deposition of fibrillar ECM components, such as fibronectin and collagen ^5^. However, brain ECM is composed primarily of proteo- and glycosamino-glycans and is largely devoid of fibrillar proteins ^30^. We, therefore, used a panel of lectins and reagents recognising carbohydrate moieties (such as hyaluronic acid-binding protein (HABP) and anti-chondroitin sulphate) to screen for EV-driven alterations to glycan/polysaccharide species in ECM deposited by astrocytes. Most carbohydrate species were not detectably different between the ECM deposited by naïve and EV-educated astrocytes (not shown). However, pre-treatment of astrocytes with EVs from E2 cells drove a significant increase in the hyaluronic acid (HA) (detected by HABP) (Figure 5A), and a moderate, but not significant, decrease the chondroitin sulphate (CS) (Figure S3A) content of the ECM they deposited. Conversely, pre-treatment of astrocytes with EVs from the less invasive G7 GBM cells, or EVs from E2 cells in which either mutant p53^273H^, PODXL or Rab35 had been deleted using CRISPR, were ineffective in this regard (Figure 5B). Furthermore, CRISPR-mediated deletion of p53^R273H^ in U373 cells significantly reduced the HA content of ECM deposited by astrocytes treated with EVs from this GBM cell line (Figure. 5B). HA has a well-established role in promoting migratory and invasive behaviour of many cell types ^31^ – including tumour cells – and examination of Z-projections of 3D confocal image stacks indicated that EVs from E2 cells most significantly increased the HA content of the ECM on upper face of the astrocyte cultures where it is ideally placed to interact with GBM cells plated onto it (Figure S3B). We, therefore, allowed astrocytes to deposit ECM in the presence of hyaluronidase (Hase), which catalyses hydrolysis of HA, and this led to the deposition of ECM with reduced HA content (Figure 5C). We next plated GBM cells onto ECM generated in the presence and absence of Hase. This indicated that the ability of EV-pre-treated astrocytes to generate ECM which supports GBM cell migration was completely ablated by Hase treatment, whilst Hase did not alter the (limited) ability of ECM deposited by EV-naïve astrocytes to support GBM cell migration (Figure 5D).

**Figure 5:**
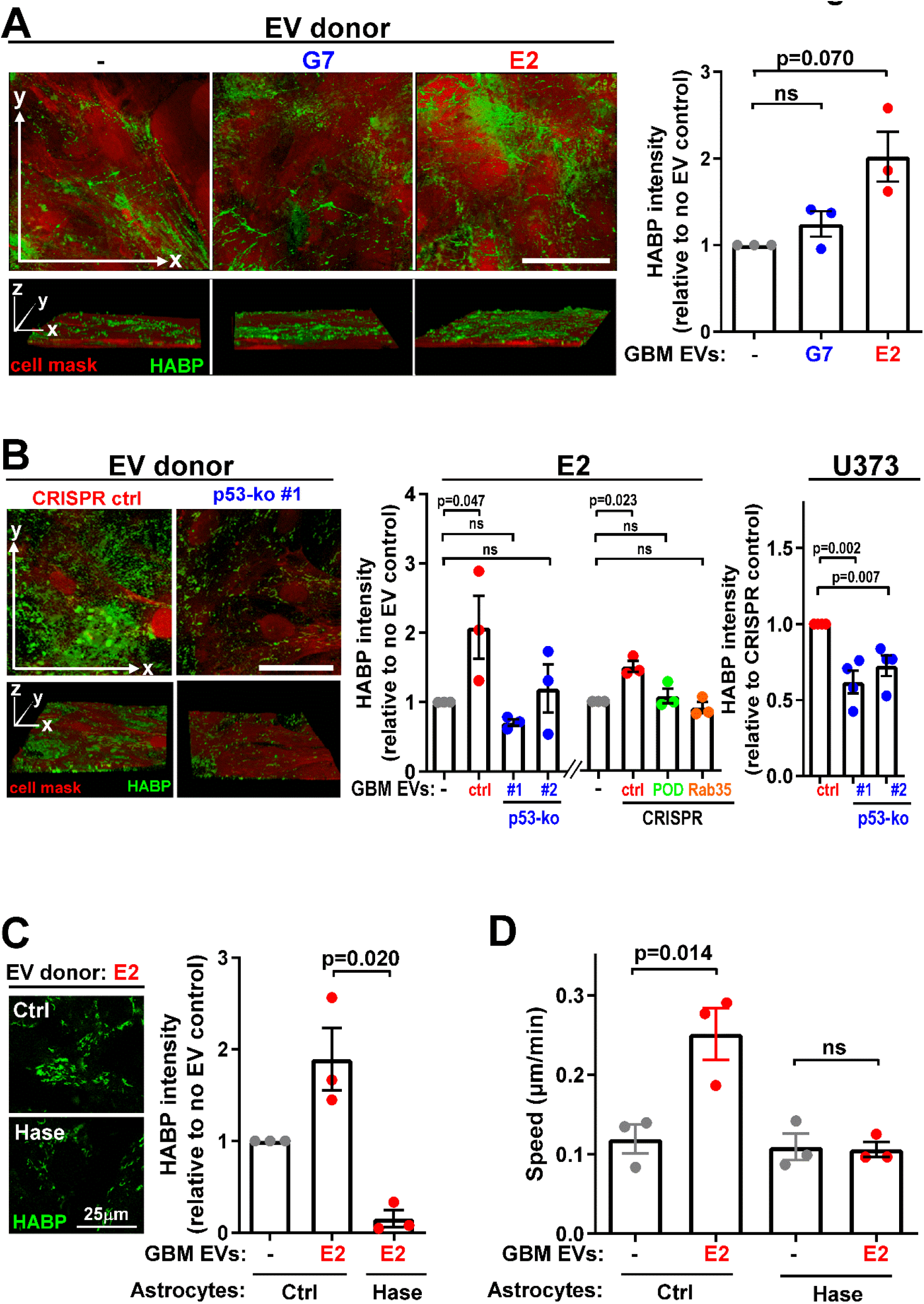
EVs from GBM cells promote GBM migration by influencing HA content of astrocyte ECM. **(A, B)** Astrocytes were incubated with EVs from G7 or E2 GBM cells (A), and E2 or U373 cells in which p53, PODXL or Rab35 had been deleted using CRISPR/Cas9 as indicated (B). Astrocytes were then allowed to deposit ECM for 6 days, stained with HABP (green) and cell mask (red) and imaged using fluorescence confocal microscopy; x/y (upper panels) and x/y/z (lower panels) projections are displayed. Bars 50 μm. HABP was quantified using Image J. Values are mean ± sem (n=3 independent experiments, paired t-test; 7 fields/condition/experiment. **(C, D)** Astrocytes were incubated with EVs from E2 GBM cells and then allowed to deposit ECM for 6 days in the presence or absence of type I-S Hase (50 µg/mL). HA content of the ECM was assessed as for (A) **(C)**, and migration of E2 cells on Ctrl and Hase-treated ECM was determined as for Fig. 1C **(D)**. Values are mean ± sem (n=3 independent experiments, paired t-test, 60 cell-tracks/condition/experiment)

These data indicate that GBM cells expressing the p53^273H^ mutant release EVs which, by virtue of their PODXL content, encourage astrocytes to deposit ECM with increased HA content which, in turn, promotes GBM cell migration.

### PODXL drives mutant p53^273H^-driven infiltrative behaviour of GBM *in vivo*

Having established three key components – mutant p53^273H^, PODXL and Rab35 – necessary for release of EVs which control astrocyte ECM deposition, and described mechanistic details of how they achieve this, we wished to determine whether these components influence GBM infiltration and invasiveness *in vivo*. We injected mutant p53, PODXL or Rab35 knockout E2 cells (or CRISPR-control) into the right striatum of CD1 nude mice. 9 weeks after injection the number of (proliferating) tumour cells in coronal brain sections (2 consecutive, 50μm sections per mouse) was quantified by Ki67-staining followed by automated cell counting. We have found that Ki67 staining clearly identifies the nuclei of tumour cells in a way that enables automated analysis. However, Ki67 staining may, due to its inability to detect non-proliferating cells, underestimate the number of invading tumour cells. We, therefore, confirmed that results from Ki67 and human-specific Leukocyte Antigen (to identify the human glioma cells) staining correlated closely ^16^. Moreover, automated counting was undertaken using parameters which excluded Ki67-positive resident brain cells in the subventricular zone. Knockout of mutant p53 or Rab35 reduced tumour growth to an extent which precluded us from determining how these potentially pro-invasive factors might influence infiltration (Figure S4). However, as the total number of Ki67-positive GBM cells in the brain was not significantly altered by PODXL knockout (as determined by quantification of either 2 or 5 consecutive 50 μm sections per mouse) (Figure S4), we were able to determine how PODXL expression influenced the proportion of GBM cells that had migrated from the ipsilateral to the contralateral (left) hemisphere. This revealed that loss of PODXL significantly reduced the proportion of Ki67-positive GBM cells which moved from the left to the right side of the brain, indicating that PODXL expression contributes to efficient long-range infiltrative/migration of GBM cells (Figure 6A, B).

**Figure 6:**
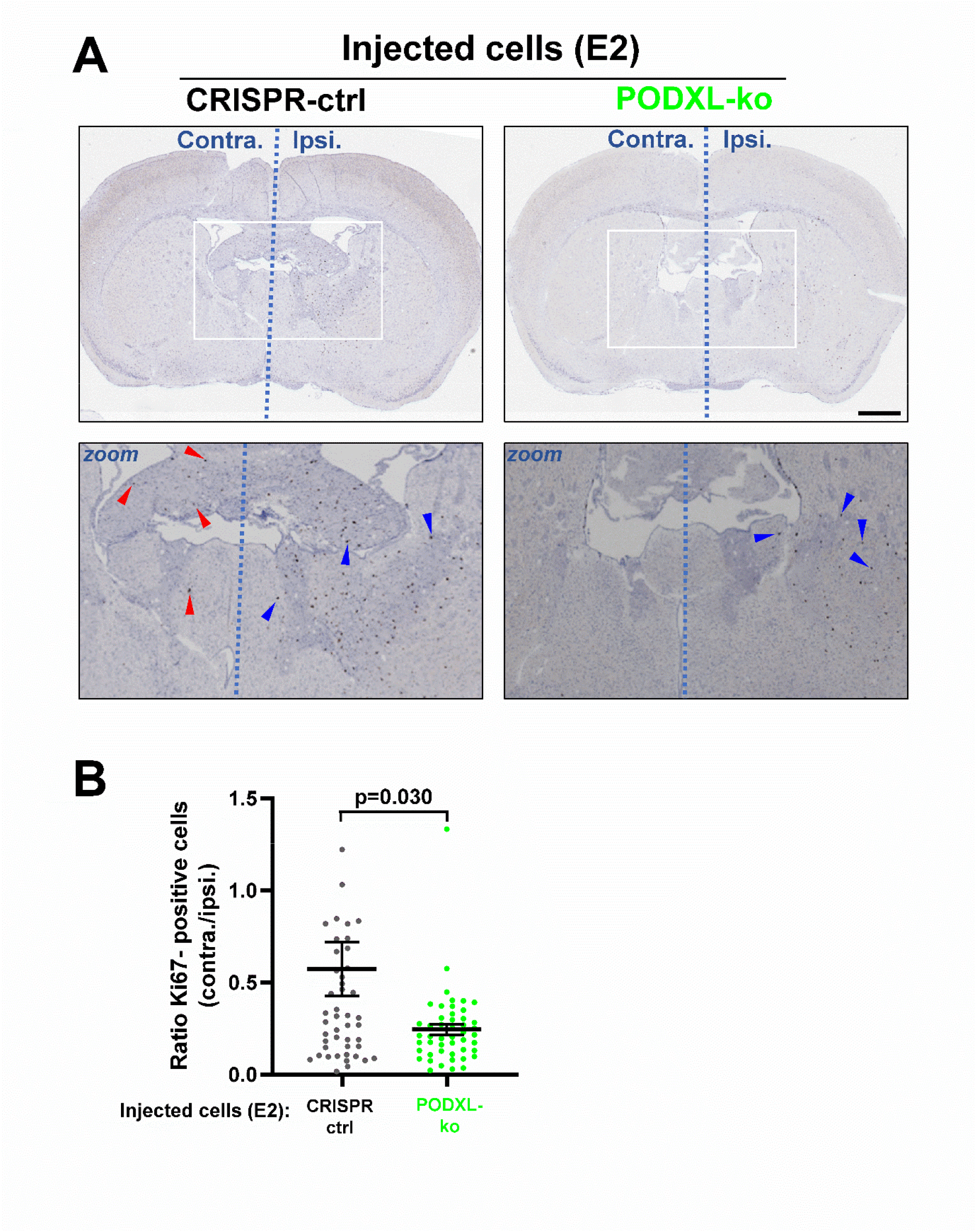
PODXL-expression is required for long-range infiltration of GBM cells. CRISPR control (CRISPR-ctrl) or PODXL knockout (PODXL-ko) GBM cells were injected into the right (ipsilateral) striatum of CD1 nude mice. 9 weeks following injection, brains were fixed and the distribution of tumour cells between ipsilateral (Ipsi.) and contralateral (Contra.) determined using Ki67 staining followed by automated image analysis. Cells remaining in the ipsilateral hemisphere and those that have migrated to the contralateral hemisphere are highlighted with blue and red arrows respectively (A). Ki67-positive cells in the contralateral hemisphere is expressed as a ratio of those in the ipsilateral hemisphere (B). Bars are mean ± sem. n=10 mice per condition. Statistic is unpaired t-test, Welch’s correction. Bar, 1 mm.

## DISCUSSION

Communication between cancer cells in the primary tumour and ECM-depositing fibroblasts in metastatic niches is central to metastasis of carcinoma ^4,5^ and, in this study, we demonstrate that a process analogous to carcinoma metastatic niche priming may occur as GBM infiltrate the brain (Figure S5). Indeed, patient-derived glioma cells can display markedly different migratory behaviour within the brain microenvironment, but this owes less to their intrinsic migratory capacity than it does to their influence on ECM deposition by glial cells. Principal amongst the similarities between this mechanism of glioma infiltration and metastatic niche priming by carcinomas is reliance on an EV-transduced signal emanating from the primary tumour cell which influences the behaviour of ECM-depositing cells. This signal is switched-on in both carcinoma and GBM cells by acquisition of a particular p53 mutation - p53^273H^ - which acts in combination with Rab35 to tune the PODXL content of tumour cell EVs into a range that influences ECM deposition by recipient cells – be they astrocytes or fibroblasts. Given the novel gain-of-function that we here describe for p53^273H^, it is interesting to consider the impact of this particular p53 mutant on glioma progression. This is especially important in light of the invasive nature of both high- and low-grade glioma which prevents curative resection. Indeed, while supratotal resection techniques significantly improve progression-free survival in low grade glioma, to date no curative treatment is available^32^. However, any extension of time to recurrence potentially provides a therapeutic window, where inhibiting further invasion of the brain by modulating the microenvironment might facilitate future treatment such as second surgery or irradiation, thus potentially enhancing patient survival. Therefore, given our evidence that p53^R273H^ can engender a pro-invasive brain microenvironment, it was important to determine whether expression of p53^273H^ might be associated with poor clinical outcomes. Examination of the merged Cell 2016 LGG and GBM dataset using cBioportal indicated that 40% tumours (319/794) displayed mutations of the p53 gene, with the R273H locus being by far the most common p53 mutation – accounting for 22% (69/319) of all p53 mutations (Figure S6A). However, the different prognosis of the glioma subtype represented within this dataset prevented us from drawing general conclusions regarding survival. As the database also lacks important information dictating the prognosis of GBM such as tumour location, resection status and adjuvant treatment, we turned to the TCGA LGG PanCancer Atlas dataset ^33^ to determine whether IDH-mutant 1p/19q non-codeleted astrocytoma harbouring p53^273H^ might progress more quickly than those bearing other p53 mutations. This indicated that the presence of a p53^273H^ mutation significantly reduced overall median survival by comparison with tumours bearing other p53 mutations (Figure S6B). Thus, we submit that the ability of p53^273H^ to influence the PODXL content of EVs and, thereby, engender a pro-invasive brain microenvironment, may contribute to the particularly fast progression and poor prognosis of gliomas bearing this p53 mutant.

Fibroblasts respond to PODXL-containing EVs by upregulating DGKα-dependent integrin recycling; DGKα operates by generating phosphatidic acid in the plasma membrane to enable docking and fusion of endocytic recycling vesicles ^22^. Increased DGKα-dependent integrin recycling then changes alignment of fibrillar ECM deposited by fibroblasts to support tumour cell migration ^5^. Unlike fibroblasts, glial cells do not deposit fibrillar ECM so it is important to consider how DGKα-mediated vesicle trafficking might influence HA deposition by astrocytes. HA synthases are transmembrane proteins which must be plasma membrane-localised to coordinate the export and growth of HA chains on the cell surface ^34^. HA synthases, like integrins, cycle between endosomes and the plasma membrane ^35^, and it will be interesting to determine whether EVs from p53^273H^-expressing glioma influence recycling of these enzymes. Secondly, ECM-associated HA is turned-over by internalisation via endocytic (clathrin and non-clathrin dependent) and/or macropinocytic routes followed by hyaluronidase-mediated degradation in lysosomes ^34^. In addition to receptor recycling, DKGα also regulates micropinocytosis and endocytosis ^22,36,37^, and it will be interesting to determine whether EVs from glioma influence HA deposition by controlling these internalisation mechanisms. Increased HA production has been linked to GBM aggressiveness, via enhanced cancer cell proliferation, chemo- and radio-resistance and invasiveness ^38,39^. This has been attributed to engagement of HA receptors, such as CD44, on tumour cells, and it is likely that the migratory response of GBM cells to HA-rich, astrocyte-deposited ECM is mediated via CD44. HA’s anti-adhesive properties ^40^ may also contribute to increased cell migration.

Mobilisation of neutrophils and macrophages is key to carcinoma dissemination and metastasis and this, in part, is owing to the immunosuppressive microenvironments that these cells engender. Immunosuppressive myeloid populations, particularly tumour-associated macrophages, also drive glioma progression and therapy-resistance. Indeed, interaction of glioblastoma with surrounding astrocytes to induce an immunosuppressive environment has been linked to tumour progression ^41,42^. Interestingly, HA can re-educate tumour-associated macrophages to a tumour-supporting phenotype ^43,44^, and this may promote GBM infiltration. Consistently, studies in which GBM and peripheral blood cells were co-cultured indicated that glioblastoma cells can surround themselves with a halo of HA-rich glycosaminoglycans ^45^. Therefore, increased HA expression by surrounding astrocytes might serve as an additional barrier to invading immune cells while simultaneously promoting glioblastoma invasion.

It is possible that HA may influence glioma infiltration and therapy-resistance by promoting tumour cell stemness. HA in the nervous system has its peak during brain development with a subsequent decline to adulthood ^46^. In adult mice, HA is only abundant in the subventricular zone and rostral migratory stream, which are thought to constitute neural stem cell niches ^47^. Therefore, by increasing astrocyte HA secretion, mp53-expressing GBM may expand the number of stem cell niches in the adult brain, thus supporting a less proliferative stem-like phenotype capable of evading current standard treatments such as radiotherapy and chemotherapy while simultaneously increasing their infiltration to evade surgical resection.

In conclusion, this study offers a mechanistic explanation for the particularly poor prognosis of GBM which express the p53^273H^ mutant and highlights potential druggable targets both within the tumour (Rab35, PODXL) and astrocytes (DGKα) which influence the ECM microenvironment (Figure S5). Moreover, our finding that increased HA deposition by astrocytes is key to glioma cell migration will prompt further investigation into the impact of this ECM component on the immune microenvironment in the brain and the regulation of tumour stem cell maintenance and resistance to therapy.

## ACKNOWLEDGEMENTS

Many thanks to Colin Nixon and the CRUK-Beatson Institute’s histology lab for the sectioning and staining of brain slices.

---

**Supplementary table S1:** p53 mutations identified by RNAseq in the E2 and G7 patientderived glioma stem-like cell lines.

| Location | change | frequency | function of region affected | name of mutation (IARC TP53 database https://p53.iarc.fr/) | effect of mutation according to IARC database |
| --- | --- | --- | --- | --- | --- |
| 7577120 | C>T | 83% | Coding | p.R273H | deleterious |
| 7577674 | A>T | 29% | UTR | not found |  |
| 7577679 | Indel 1-2bp | ? | UTR | not found |  |
| 7578712 | Indel 1-3bp | ? | Intron | not found |  |

**B** *p53 mutations G7 cell line*
| Location | change | frequency | function of region affected | name of mutation (IARC TP53 database https://p53.iarc.fr/) | effect of mutation according to IARC database |
| --- | --- | --- | --- | --- | --- |
| 7577094 | G>A | 49% | Coding | p.R248Q | deleterious |
| 7577538 | C>T | 51% | Coding | p.R282W | deleterious |
| 7577674 | A>T | 28% | UTR | not found |  |
| 7578115 | T>C | 50% | UTR | not found |  |
| 7578645 | C>T | 52% | Intron | not found |  |
| 7578712 | Indel 2-5bp | ? | Intron | not found |  |
| 7578837 | A>G | 69% | UTR | not found |  |
| 7579472 | G>C | 49% | Coding | p72r |  |
| 7579801 | G>C | 39% | Intron | not found |  |

**Figure S1:**
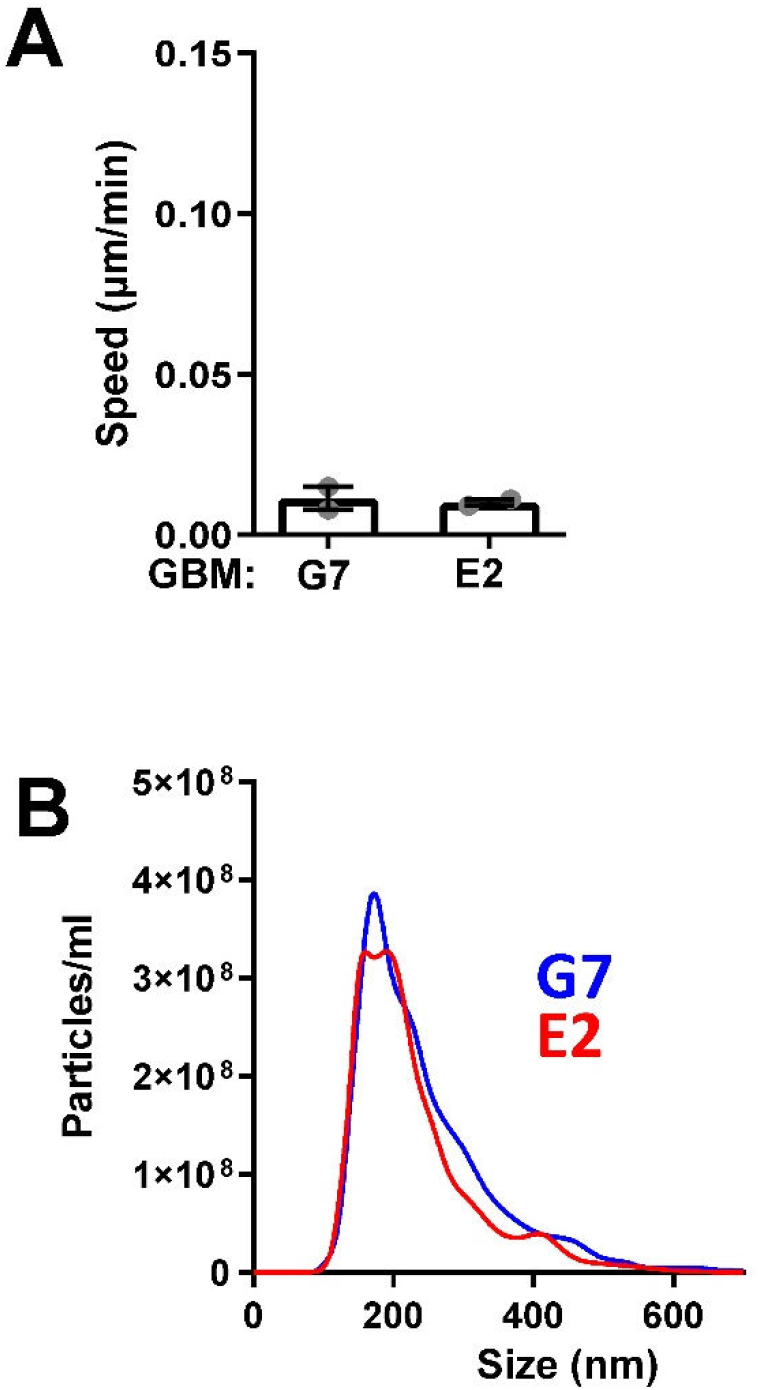
**(A) G7 and E2 GBM stem-like cells display similar migratory behaviour when plated onto mouse brain slices** GFP-expressing GBM (E2 and G7) cells were plated onto coronal mouse brain slices and their migration speed determined using time-lapse fluorescence microscopy. Values are mean ± sem (n=2 independent experiments, 60 cell-tracks/condition/experiment). **(B) Characterisation of EVs from G7 and E2 GBM cells** E2 (red) or G7 (blue) GBM cells were incubated in EV-free medium for 48 hr. EVs were then purified from conditioned medium using differential centrifugation. The number and size-distribution of EVs was analysed using nanoparticle tracking. Each line represents the mean of three independent experiments.

**Figure S2:**
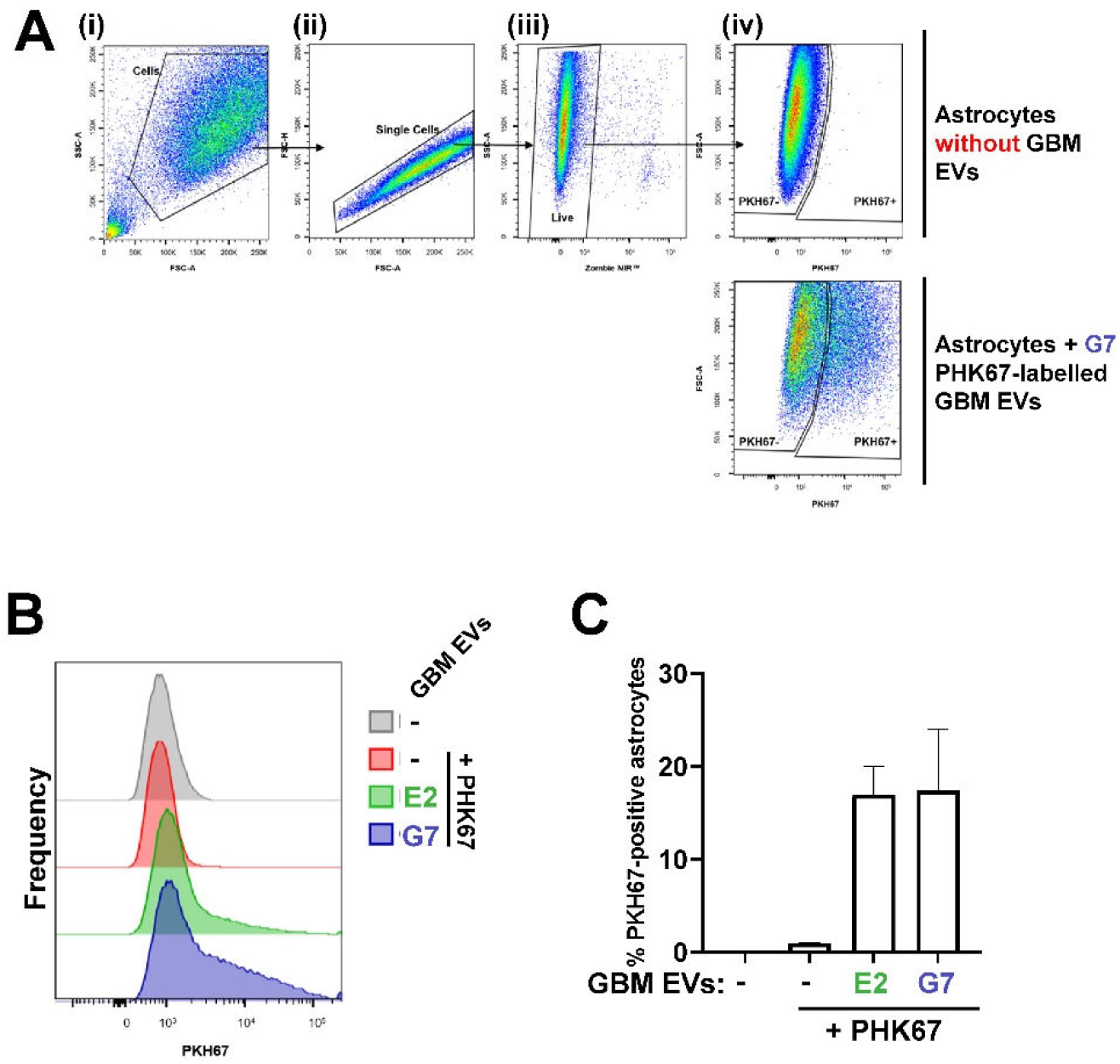
**Uptake of labelled EVs from G7 and E2 cells by astrocytes** E2 (green) or G7 (blue) GBM cells were incubated in EV-free medium for 48 hr. EVs were then purified from conditioned medium using differential centrifugation, and then labelled by incubation with PHK67 (2μM) for 5 min. Excess dye was removed by Ultracentrifugation (100,000 g for 70 min) and labelled EVs were added to primary cultured astrocytes for 24hr. Recipient astrocytes were then analysed using flow cytometry to determine uptake of labelled EVs. Flow cytometry was sequentially gated to include cells (**A**; (i)), single cells (**A**; (ii)) and live cells (**A**; (iii)), then PHK67 fluorescence was detected using an emission filter with 500 – 560 nm cut off (**A**; (iv)). The distribution of fluorescence intensities **(B)** and the proportion of recipient astrocytes which had taken-up PHK67-labelled EVs **(C)** from E2 (green) or G7 (blue) GBM cells was then determined. Values are mean ± SEM, n=6.

**Figure S3:**
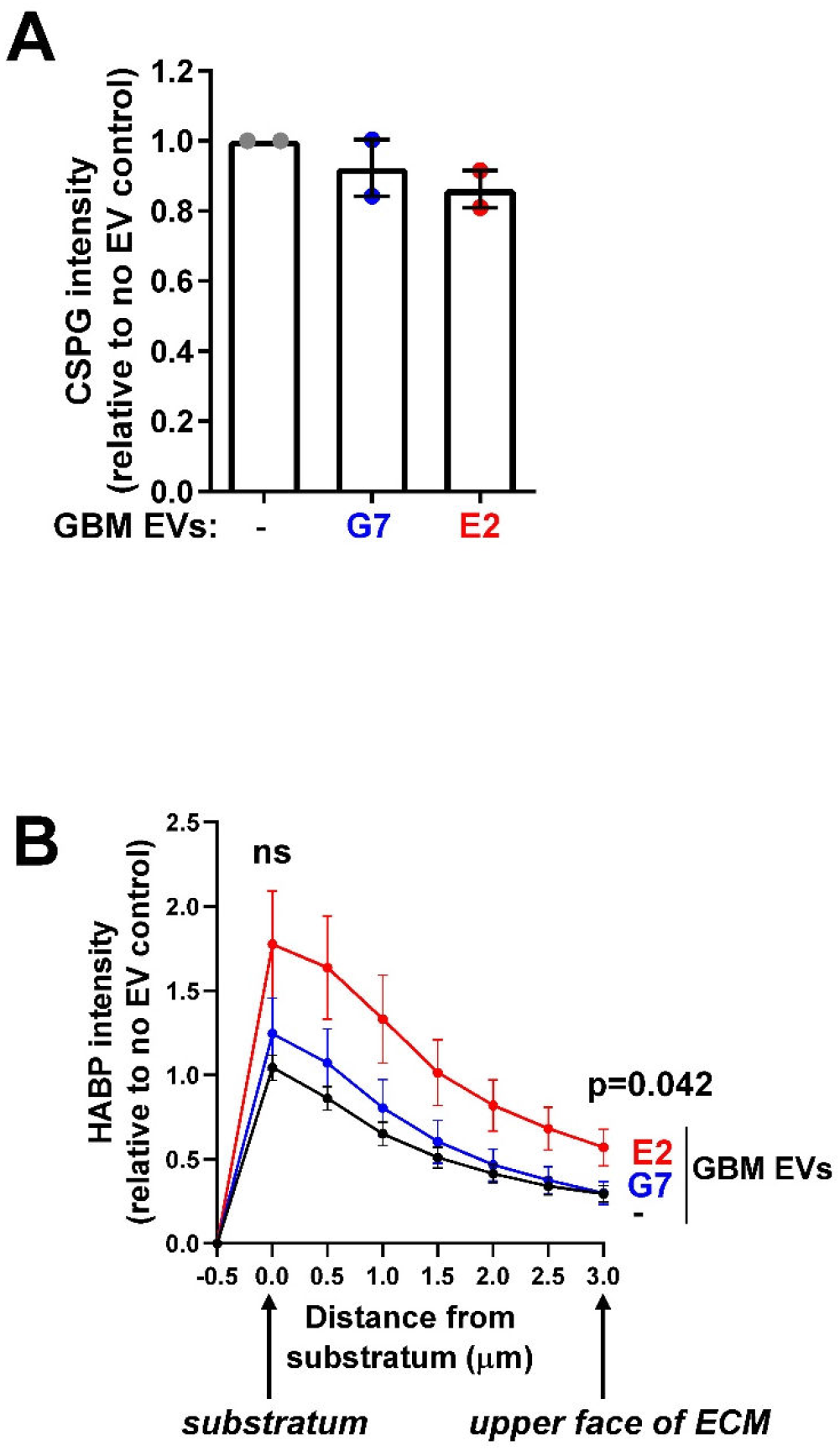
**(A) EVs from highly infiltrative E2 cells do not influence the CSPG content of brain slices** Astrocytes were incubated with EVs from G7 or E2 GBM cells. Astrocytes were then allowed to deposit ECM for 6 days, stained with an antibody recognising CSPG and imaged using fluorescence confocal microscopy. Fluorescence was quantified using Image J. Values are mean ± sem (n=2 independent experiments, paired t-test; 7 fields/condition/experiment). **(B) EVs from highly infiltrative E2 cells increase the HA content at the upper face of astrocyte-deposited ECM** Astrocytes-deposited ECM was generated as for (A) and stained with biotinylated HABP followed by fluorescent streptavidin to visualize HA. The ECM was imaged using fluorescence confocal microscopy and the quantity of HA present in optical slices at the indicated distances from the substratum was determined using Image J. The upper face of the ECM is 3.0 μm from the substratum. Values are mean ± sem of three independent experiments, unpaired t-test E2 versus G7.

**Figure S4:**
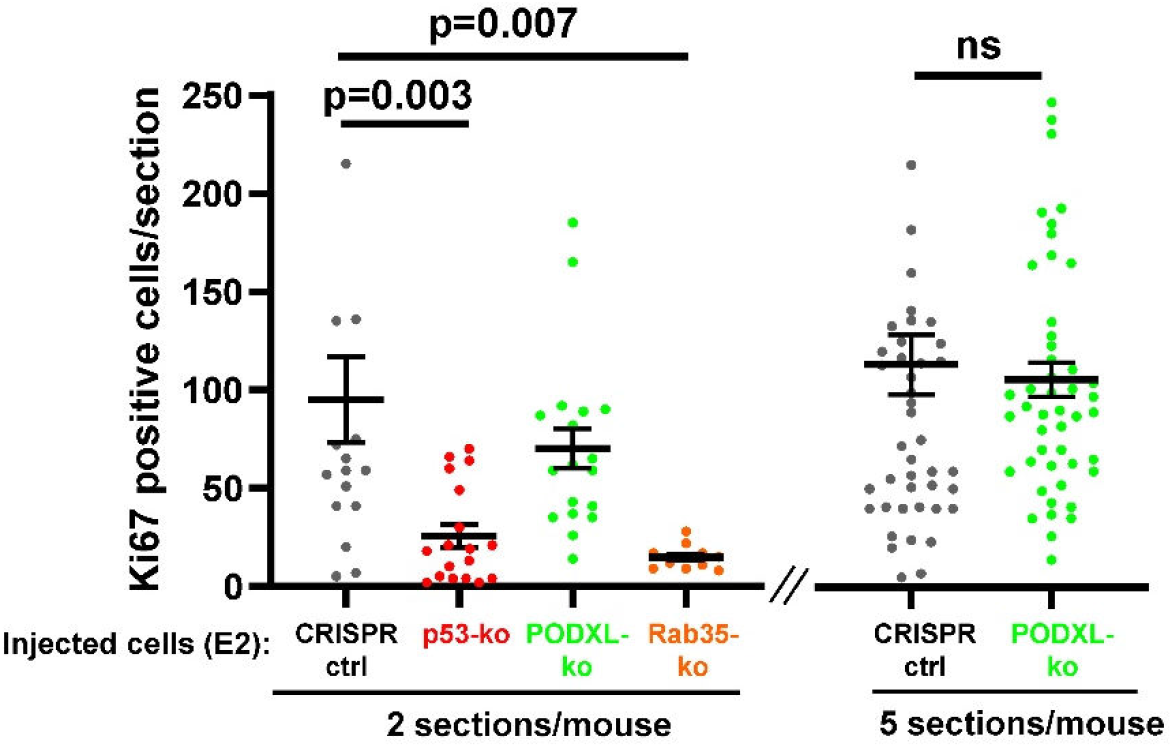
Knockout of mp53^273H^ or Rab35 opposes growth of GBM in vivo. CRISPR control (CRISPR-ctrl) GBM (E2) cells, or those in which mp53^273H^ (p53-ko), PODXL (PODXL-ko) or Rab35 (Rab35-ko) had been knocked out were injected into the right striatum of CD1 nude mice. 9 weeks following injection, brains were fixed, cut into 50 μm sections and the quantity of tumour cells present in the section determined by staining for Ki67 followed by automated image analysis. The left- and right-hand graph represent data from 2 and 5 consecutive, 50 μm sections per mouse. Bars are mean ± sem. The dots represent the number of Ki67-positive cells present in each individual brain slice. Statistic is unpaired t-test with Welch’s correction, ns is not significant.

**Figure S5:**
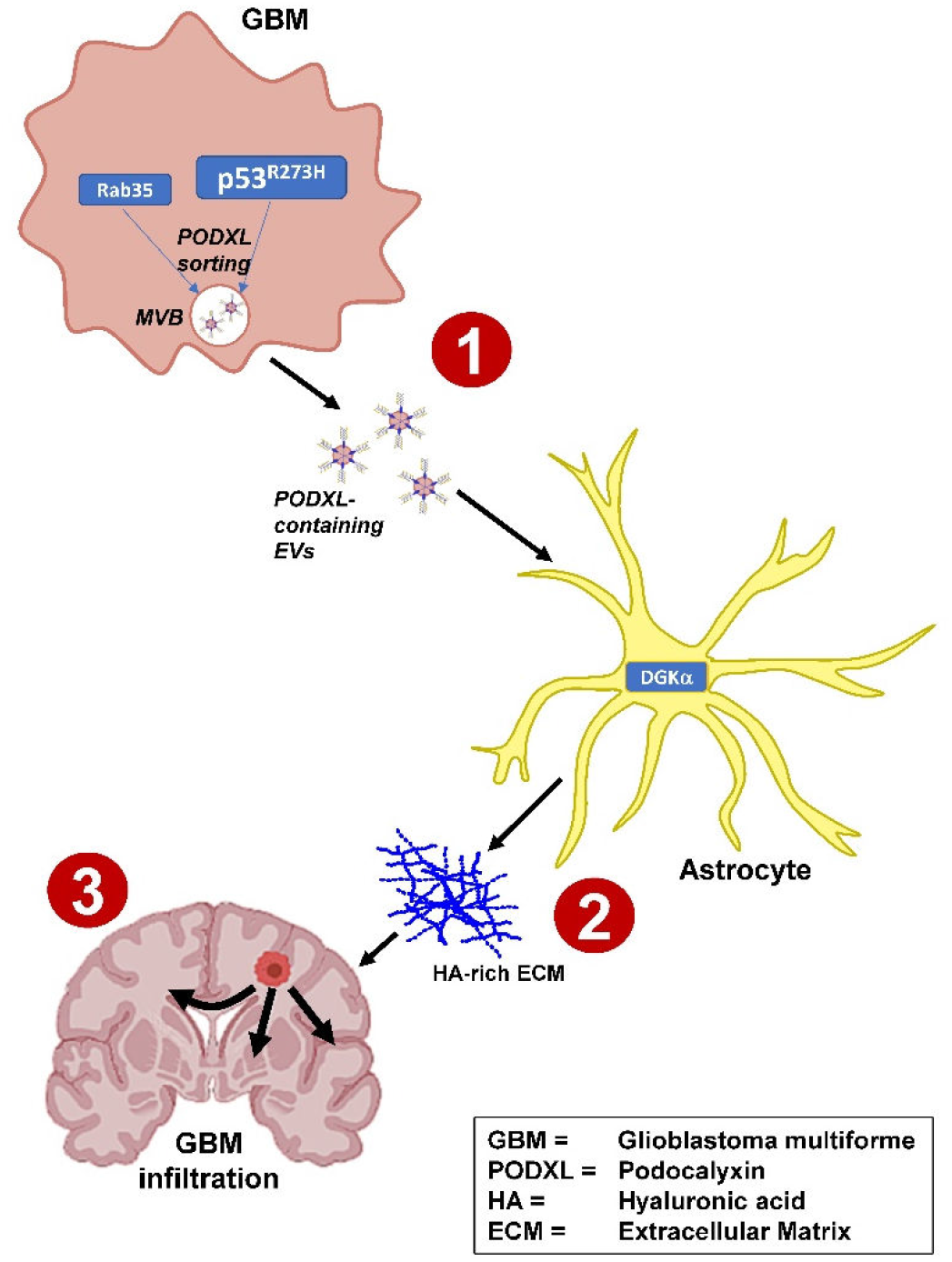
Schematic summary. **[1]** In glioblastoma cells, a gain-of-function p53 mutation acts in combination with a GTPase of the Rab family (Rab35) to control sorting of a sialomycin, called podocalyxin, into small extracellular vesicles (EVs). **[2]** Podocalyxin containing EVs from glioblastoma cells act on astrocytes to influence the type of extracellular matrix that they produce. They encourage astrocytes to deposit an extracellular matrix which is particularly rich in the glycan, hyaluronic acid (HA). **[3]** HA-rich extracellular matrix, in turn, encourages the glioblastoma cells to be more invasive and to migrate long distances to infiltrate the brain.

**Figure S6:**
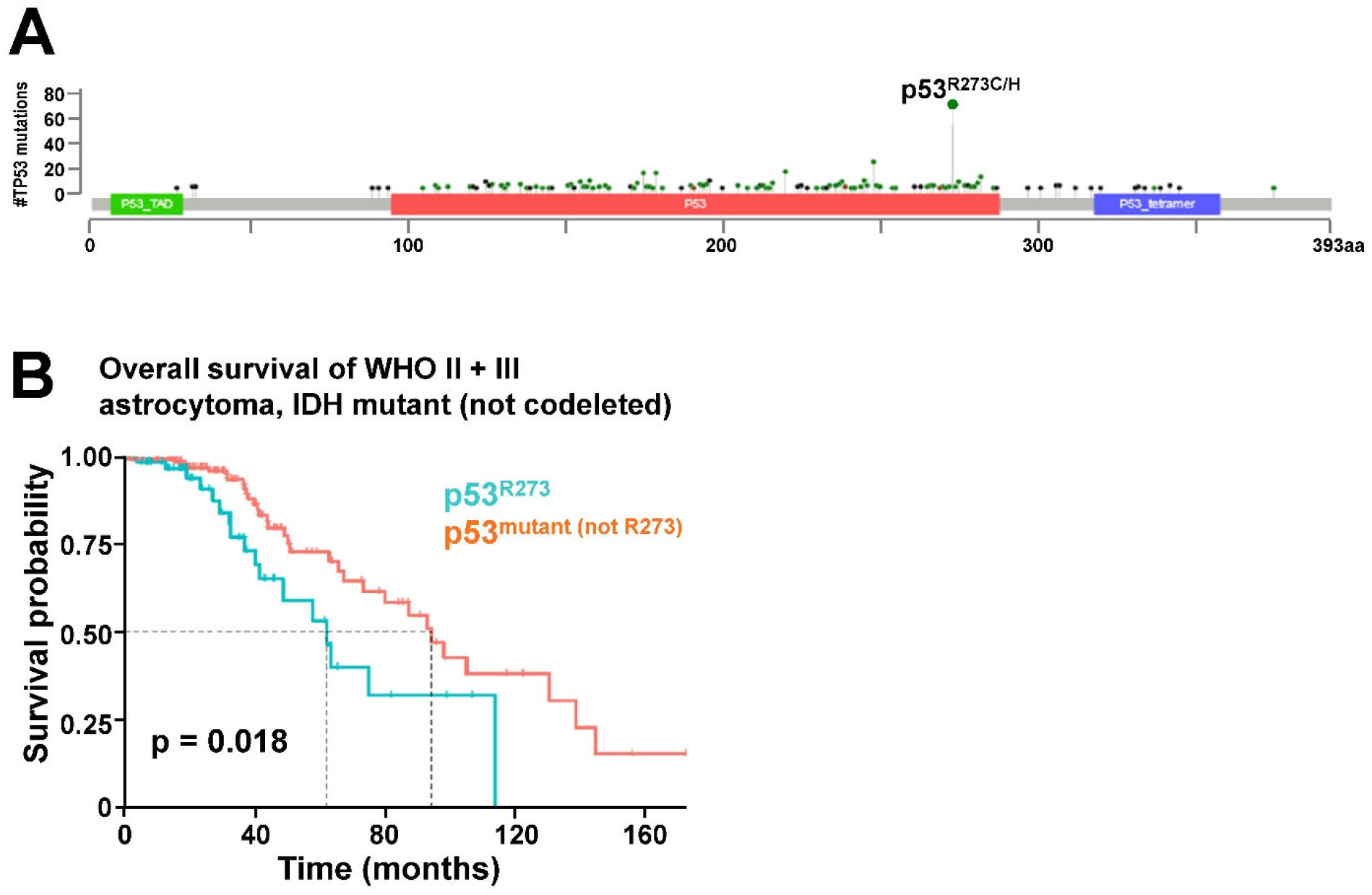
p53 mutations in GBM. **(A)** Distribution of p53 mutations within a merged TCGA cohort of LGG and GBM. The R273 locus, is by far the most common mutation across this panel of different tumours. **(B)** Comparison of the overall survival of patients bearing p53^273H^ -expressing WHO II + III IDH mutant (not codeleted) astrocytoma (blue line) with patients bearing tumours of the same subtype, but that contained p53 mutations other than p53^R273H^ (orange line).

